# Control of Molecular Biochemistry and Cell Injury Responses through Highly Ordered Supramolecular Assembly of Flavonoids

**DOI:** 10.1101/2024.04.03.587976

**Authors:** Charles B. Reilly, Sylvie G. Bernier, Sanjid Shahriar, Viktor Horvath, Michael Lewandowski, Emilia Javorsky, Bogdan Budnik, Donald E. Ingber

## Abstract

Flavonoids are phytonutrients commonly found in plant-based foods and are generally known for their health benefits. However, their utility as potential therapeutics has not been explored because their presence in drug development tests can lead to false positives due to non-specific binding. Here, we employed molecular dynamic simulations (MDS) to examine flavonoid behavior and discovered that they form highly organized supramolecular assemblies that physically interact with disordered regions of enzymatic proteins and can physically interlink multiple protein molecules. These flavonoid assemblies adopt secondary structural patterns like those found in proteins and nucleic acids, and they physically influence molecular movement and tertiary protein structure, thereby modulating the biochemical activities of a diverse range of enzymes. Moreover, in the presence of flavonoids, human cells are protected against injury caused by ultraviolet radiation. These findings unveil a novel form of biochemical regulation wherein small molecules can modulate the function of larger proteins by forming supramolecular assemblies which results in enhanced molecular and cellular resilience.

**Single Sentence Summary:** Molecular dynamic simulations led to the discovery that flavonoid phytonutrients can self-assemble into highly ordered supramolecular structures that interact with enzymatic proteins, slow biochemical activities, and protect cells against injury.

## Introduction

Flavonoids are a subclass of polyphenols found in many fruits, vegetables, nuts, seeds, and other plant-based foods. They have many potential health benefits, including anti-inflammatory and anti-cancer activities, ostensibly based on their antioxidant properties^1–5^. However, flavonoids, such as quercetin ^6–10^, are commonly excluded from high-throughput drug screening programs because they belong to a class of small molecules that can cause pan-assay interference (PAINS), leading to false positive results^11^. This activity is assumed to be due to non-specific binding of flavonoid molecules to proteins or other biological targets. Therefore, using PAINS filters^12^ in high-throughput screening has been critical to reducing false positive hits and improving overall drug screening efficiency and accuracy. This efficiency is important in conventional drug design, where the desired therapeutic behavior is based on a drug or drug-like molecule fitting into a specific binding site on a target protein.

Given the many known health benefits of flavonoids and their high oral tolerability, they could represent potential therapeutics that would be missed because they are now excluded from conventional drug discovery pipelines. Therefore, we carried out biochemical studies and molecular dynamics simulations (MDS) to better understand how flavonoids might influence the biochemical activities of multiple unrelated enzymes. In these studies, we discovered that flavonoids slow molecular biochemistry through higher-order self-assembly into multimolecular structures that physically impinge on the molecular motion of enzymatic proteins.

## Results

First, we carried out experimental studies to investigate the impact of 58 different flavonoids on biochemical activities using seven different enzyme assays, including Arginase 1 (ARG1), Lysine demethylase 4C (KDM4C), Lysozyme, MAP/microtubule affinity-regulating kinase 4 (MARK4), Histone-Lysine N-methyltransferase (NSD2), Tyrosine-protein phosphatase non-receptor type 1 (PTP1B), and Siturin-3 (SIRT3) (**Extended Data Table 1**). Results from enzyme activity analysis revealed that subtle variations in flavonoid structure led to varying specificity in enzyme activity inhibition (**Extended Data Table 2**). We focused on 3 of the 58 flavonoids that showed distinctly different enzyme specificities (quercetin, isoquercitrin, and quercitrin) to gain further insight into the underlying mechanism responsible for their observed inhibitory effects. Quercetin, the smallest compound with a core flavonoid structure, inhibited all tested enzymes, while isoquercitrin and quercitrin, which share the same core flavonoid structure as quercetin, demonstrated varied levels of inhibition (**Fig. 1a**). The primary structural difference between quercetin and the other two flavonoids is the presence of a glycosidic sugar group instead of a hydroxyl group in the number 3 position of the C-ring —glucose in isoquercitrin and rhamnose in quercitrin (**Fig. 1b**). These glycosidic groups further vary based on the position of their hydroxyl groups.

**Fig. 1:**
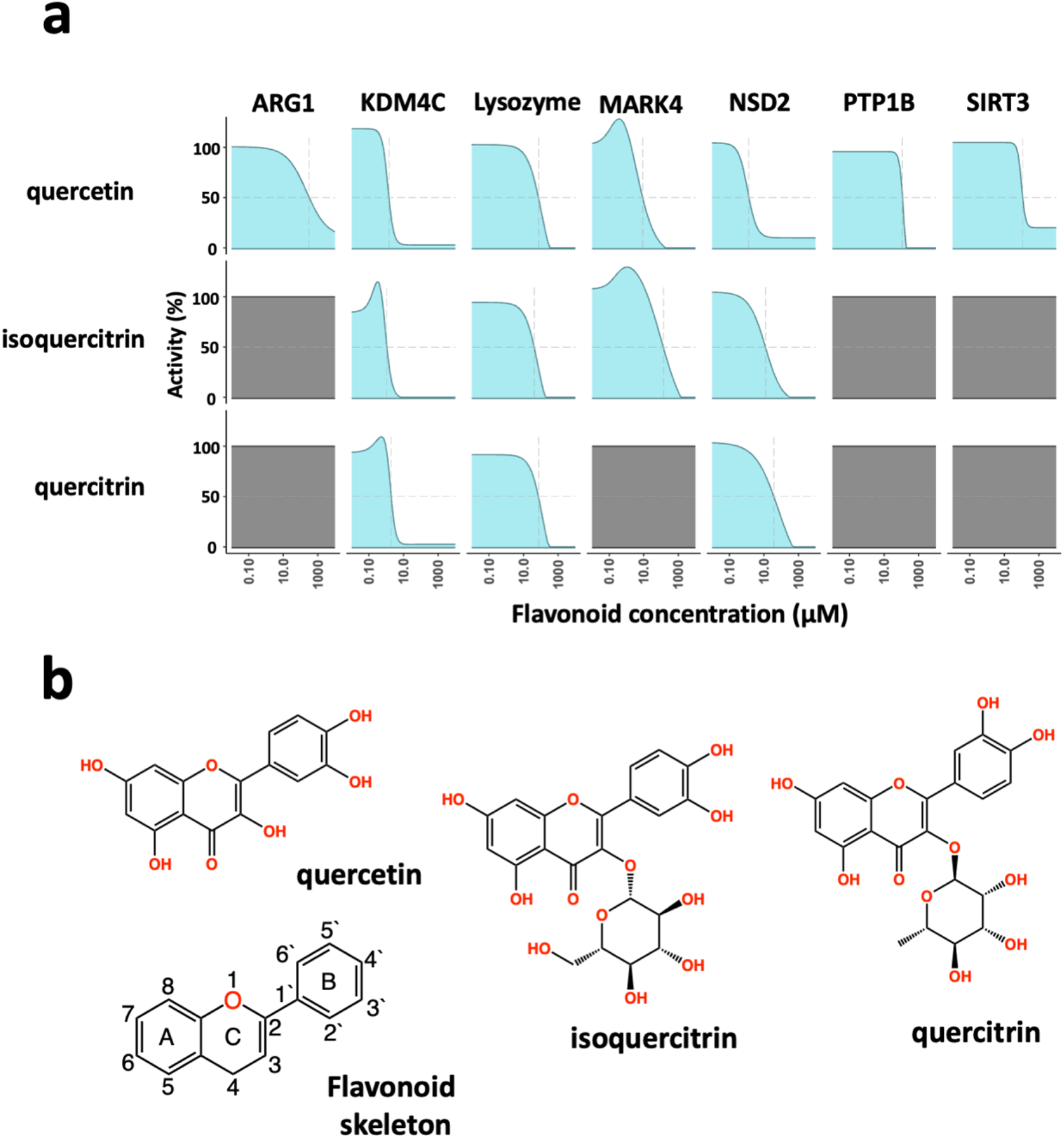
Broad-spectrum inhibition of enzymatic activity by flavonoids. **a,** Dose-dependent inhibition of enzyme activity. In biochemical assays, the flavonoids quercetin, isoquercitrin, and quercitrin show different enzyme inhibition of Arginase 1 (ARG1), Lysine demethylase 4C (KDM4C), lysozyme, MAP/microtubule affinity-regulating kinase 4 (MARK4), Histone-Lysine N-methyltransferase (NSD2), Tyrosine-protein phosphatase non-receptor type 1 (PTP1B), and Siturin-3 (SIRT3). Quercitin showed concentration-dependent inhibition of all seven enzymes; all three flavonoids blocked the enzymes KDM4C, Lysozyme, and NSD2; while quercitrin performed similarly to isoquercitrin, it did not inhibit the MARK4 enzyme. **b**, Chemical structures of the flavonoids, quercetin, isoquercitrin, and quercitrin, as well as the core flavonoid skeleton.

To gain mechanistic insight into how these subtle structural variations in flavonoids may lead to different impacts on enzyme inhibition, we conducted MDS on two enzymes: MARK4 and PTP1B, which showed varied activity in the presence of the three highlighted flavonoids. Quercetin inhibited both enzymes, quercitrin did not affect either enzyme, and isoquercitrin inhibited MARK4, but not PTP1B. The starting structures of enzymes used in the simulations were generated using AlphaFold^13^, and they both exhibited disordered regions with low structural integrity. Simulations were performed with a consistent stoichiometry (1:1 ratio by mass of enzyme protein to quercetin).

Surprisingly, 100 ns simulations revealed that the flavonoids self-assembled into highly ordered supramolecular structures (**Fig. 2a**). These flavonoid structures interacted with the enzymes differently depending on the specific flavonoid. Notably, these interactions influenced the conformational states of the disordered enzyme protein regions, and the supramolecular flavonoid structures formed in these protein regions varied based on the flavonoid type. With both MARK4 and PTP1B enzymes, the supramolecular flavonoid structures impacted the protein backbone structure near the enzyme’s active site (**Fig. 2b**). These subsequent structural changes to the disordered protein regions likely influence the substrate orientation and its access to the enzyme’s active site, suggesting a potential mechanism for enzyme regulation.

**Fig. 2:**
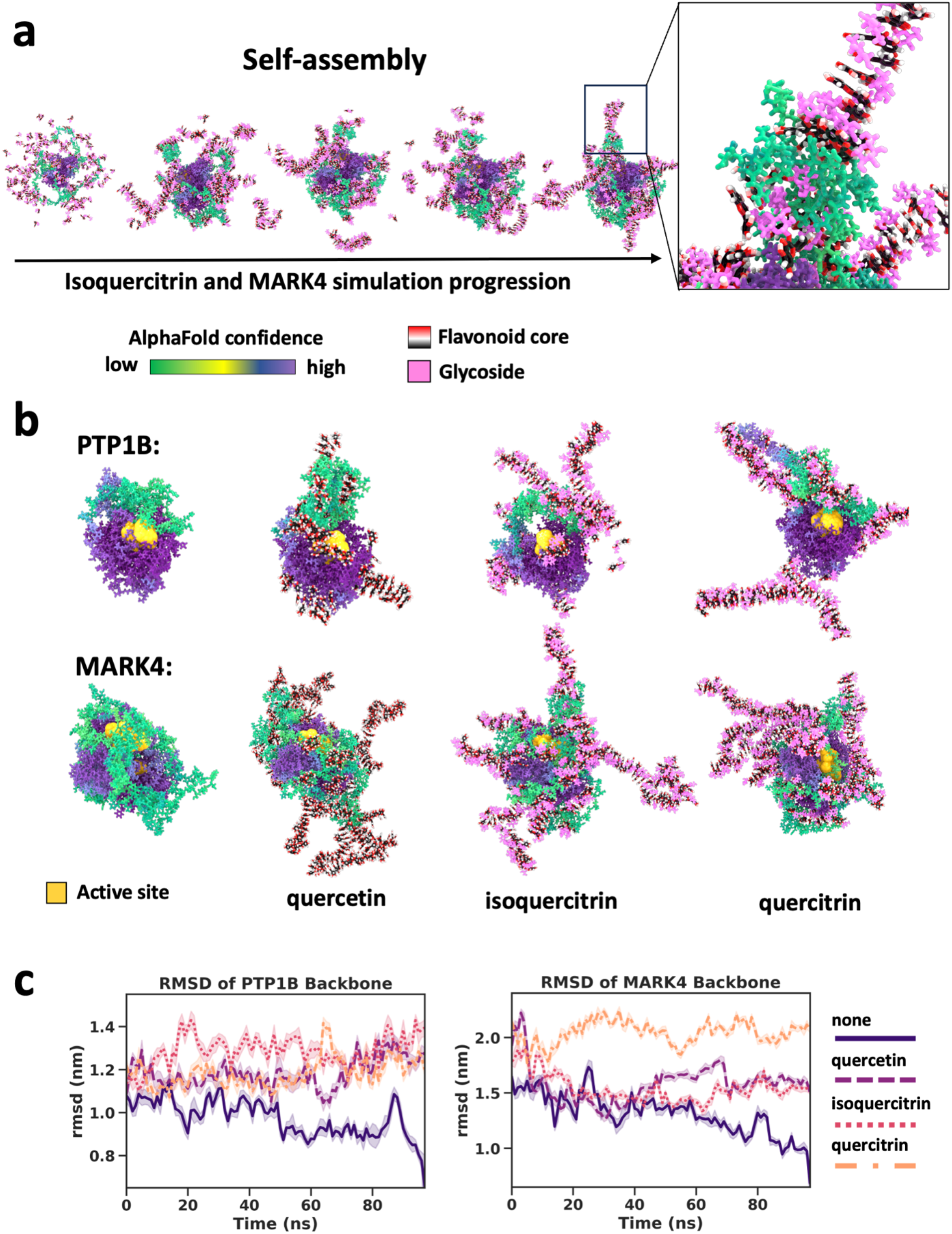
Flavonoids form higher-ordered structures that influence the molecular dynamics of proteins. **a,** States taken every 25 ns from an MD simulation of a MARK4 enzyme molecule in the presence of isoquercitrin over the course of a 100 ns MD simulation. Self-assembly of a supramolecular structure involving the flavonoids and the protein. The proteins are colored based on AlphaFold confidence score. The flavonoid cores are rendered based on element and the flavonoid’s glycosidic moiety is pink. **b,** States taken after 100 ns MD simulation of PTP1B and MARK4 enzymes in the presence of flavonoids with the active site enlarged and in yellow. (left to right) Protein alone, protein and quercetin, protein and isoquercitrin, and protein and quercitrin. **c,** RMSD analysis of simulations in panel **b**, PTP1B core enzyme backbone for residues 3 to 277 (left) and MARK4 residues 59 to 310 (right) in the presence of flavonoids. Analyses were performed with the reference state taken from the end of a 100 ns MDS of the corresponding protein backbone in the absence of flavonoids.

For the PTP1B enzyme, only quercetin showed inhibitory activity, which correlates with the crowding of the enzyme’s active site at its quercitin-stabilized regions (**Fig. 2b**). In contrast, the enzyme’s active site appeared to be less crowded when isoquercitrin and quercitrin were present, and no enzyme inhibition was observed. For the MARK4 enzyme, its active site was crowded in simulations with both quercetin and isoquercitrin, while open in the presence of quercitrin. These observations align with our assay results, where both isoquercitrin and quercetin inhibited MARK4, while quercitrin did not (**Fig. 1a**).

In addition to stabilizing the disordered enzyme protein regions and their impact on substrate access to the enzymes’ active site, we investigated whether flavonoid supramolecular assemblies alter the structural dynamics of core regions crucial for an enzyme’s catalytic activity. These regions were defined as crystallizable enzyme domains based on UniProt assignments (AC: P108031 and Q96L34 for PTP1B and MARK4, respectively), which generated high-confidence AlphaFold structures. We conducted root mean squared deviation (RMSD) analysis on the simulations using the protein backbone of residues 59 to 310 in MARK4 and 3 to 277 in PTP1B to assess protein flexibility. The RMSD values were calculated with reference to a simulation of the enzyme in the absence of flavonoids after 100 ns.

The results indicated that all three flavonoids induce structural alterations in the core regions of PTP1B compared to its reference state (**Fig. 2c**). In the case of MARK4, isoquercitrin and quercetin exhibited similar structural deviations from the reference structure, while quercitrin caused more significant variations. These findings are again consistent with the effects of these flavonoids on enzyme activity *in vitro*, where both isoquercitrin and quercetin demonstrated similar inhibitory activity against MARK4, and quercitrin had no effect. These findings suggest that quercitrin may stabilize an active state in the protein, while isoquercitrin and quercetin stabilize an inactive state. When we performed harmonic ensemble similarity (HES) analysis on the last ten ns of the simulation to gain additional insights into relative differences in dynamics following the formation of supramolecular structures (**Extended Data** Fig. 1), we found that the dynamics also varied depending on the specific flavonoid present for both enzymes.

Interestingly, the flavonoid supramolecular assemblies formed fibers. Fiber formation also has been observed in ensembles of intrinsically disordered proteins (IDPs)^14^, which can protect cells against environmental stressors. Fiber formation is crucial for proteins that make up the cytoskeleton via non-covalent assemblies, which structurally regulate cellular physiology through the transmission of mechanical forces^15,16^. Given their length, these long self-assembled flavonoid structures also could influence protein-protein interactions, extending their potential impact beyond the modulation of enzyme action.

To investigate the assembly of flavonoid supramolecular structures in the context of multiple protein molecules, we conducted MDS of flavonoid self-assembly in the presence of two Lysozyme molecules. During simulation, the enzyme molecules became interconnected by the flavonoid assembly, forming a unified structure (**Fig. 3a** and **Extended Data** Fig. 2). Analysis of the radius of gyration (RGYR) confirmed that the flavonoid assembly resulted in physical interconnections between neighboring Lysozyme molecules, as the entire unit of two protein molecules and bound flavonoid structures moved together as a single mass, which was indicated by a reduction in the variation of RGYR values (**Extended Data** Fig. 2a). Moreover, MDS analysis and visualization of force distributions within the self-assembled structures (**Extended Data** Fig. 2b,c) revealed that forces could be transmitted between the two protein molecules via the flavonoid assemblies. This mechanotransduction mechanism via allostery between different molecules is a common feature observed in protein complexes that undergo conformational shifts to fulfill their functional roles^15–18^.

**Fig. 3:**
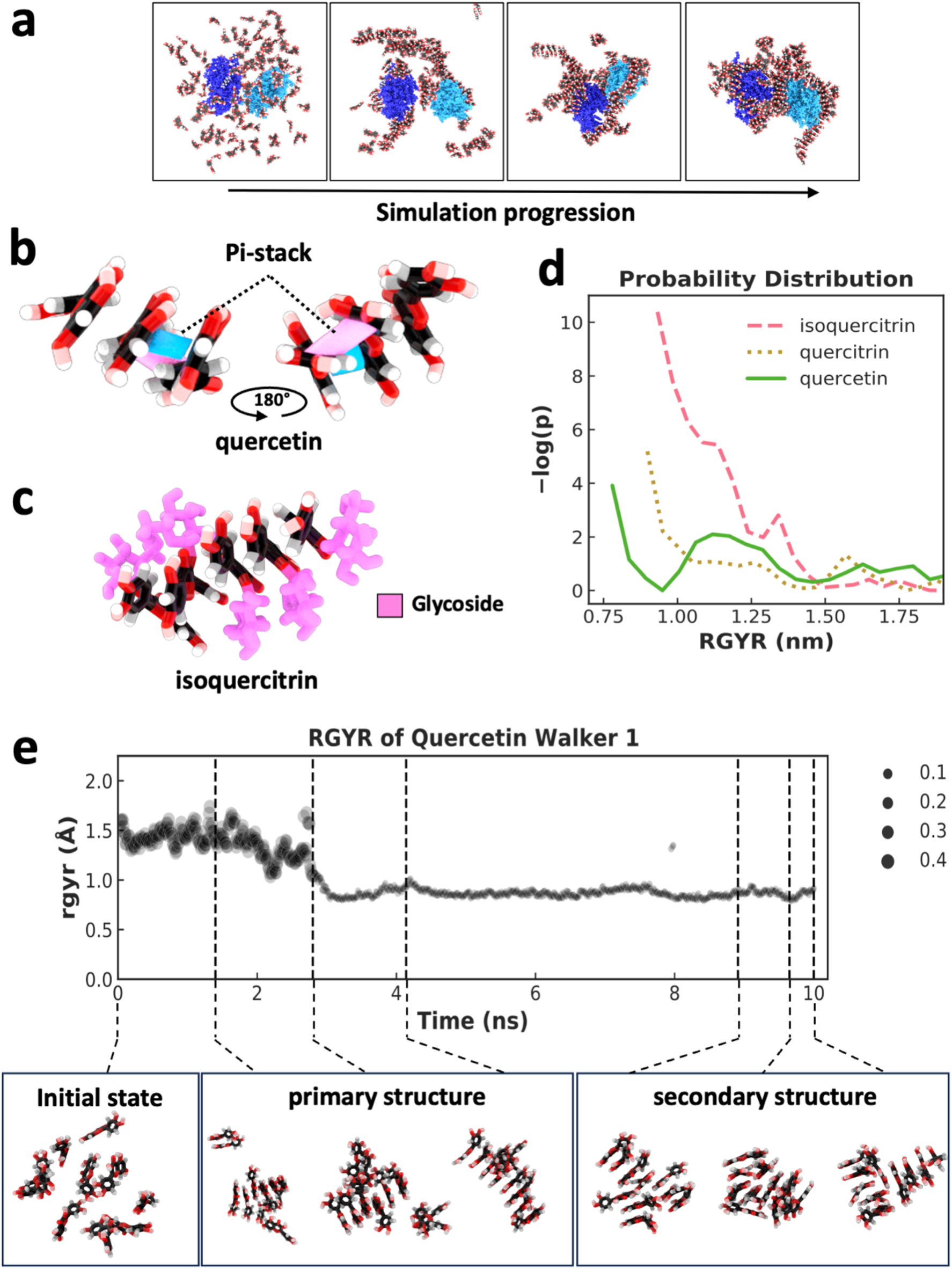
Supramolecular assembly formation and weighted ensemble simulations of flavonoids. **a,** States taken every 100 ns across a 400 ns MDS of two lysozyme molecules (dark blue and light blue) and 128 quercetin molecules, indicating the self-assembly of a supramolecular structure comprised of flavonoid and two initially separated protein molecules. **b,** Pi-stacking (shown in pink and blue between two molecules) leading to the assembly of three quercetin molecules. **c,** assembly of five isoquercitrin molecules via pi-stacking of a flavonoid core. Glycosidic groups (pink) are positioned on the outside of the assembly forming a “sugar backbone”. **d**, Indicator of assembly size and compactness using a weighted probability distribution of supramolecular structures with different RGYR. The plot represents distribution across 500 ns of simulation using 50 walkers in a weighted ensemble simulation with an RGYR distance function. **e,** RGYR trace and weighting over the 10 ns trajectory of a single walker (top) and a selection of quercetin states taken from across the trajectory (bottom) with indicated RGYR. Primary and secondary structure examples are shown along with initialization states that lack a supramolecular structure.

Through visual inspection by MDS, the supramolecular flavonoid structures that formed underwent self-assembly as a result of pi-stacking mediated by the specific flavonoid’s polyphenol substructure. MDS visualization of quercetin undergoing this process shows how the flavonoid core can form two pi-stacking interactions at two angles (**Fig. 3b**). Pi-stacking of the flavonoid core strikingly resembles the stacking of base pairs within nucleic acids. The glycosidic sugar groups in quercitrin and isoquercitrin also seemed to influence the assembly process (**Fig. 3c**). For example, when the glycoside sugar is present as it is in isoquercitrin, the supramolecular structure forms so that the sugar is on the outside of the pi-stacking, which leads to the formation of a sugar backbone and the whole structure takes on a helical form, again like in nucleic acids (**Fig. 3c**)^19^.

To quantitatively assess whether the glycosidic sugar groups promoted or hindered the formation of ordered structures, weighted ensemble simulations (WES) were conducted to explore the associated free energy landscape. A WES system was set up for each of the three flavonoids with ten molecules in explicit solvent, employing 50 walkers and a total simulation time of 500 ns using a resampler to utilize the RGYR (which measures the average distance of atoms from the center of mass or compactness) as a distance function. A free energy profile was calculated by normalizing the weighted distribution of RGYR values, followed by transformation using -ln(p) to obtain a free energy-like value ^20^, and then each flavonoid’s resulting −log(*p*) probability distribution was calculated (**Fig. 3d**). The RGYR of WES initialization (when no self-assembly has occurred) was ∼1.5 nm, and assembly caused the RGYR to decrease. The plot in **Fig. 3d** indicates significant separation in the probability distribution of the different flavonoids forming structures smaller than 1.5 nm, with quercetin showing the highest probability of developing the most compact structures (smaller RGYR) as opposed to the flavonoids with glycosidic sugar groups.

The weighted RGYR of a single representative walker lineage for a 10 ns trajectory for the quercetin system in shown in **Fig. 3e**. Seven states across the trajectory are shown (**Fig. 3e bottom**) to indicate the transition from a state with RGYR ∼1.5 nm to a state with < 1 nm and the formation of primary and secondary structures throughout the trajectory. In addition to parallel pi-stacking, there are other forms^21^ of interactions involving pi systems, and this can lead to altered geometry, which we observed in the form of primary and secondary structural forms facilitated by pi-stacking interactions and hydrogen bonding of hydroxyl groups previously identified through visualization. The primary structure of the flavonoid fiber has one stack of molecules connected by pi-stacking, and a secondary structure leads to additional dimensionality in the supramolecular structure. This secondary structure is formed when a second primary structure branches out perpendicular or parallel to the first. This secondary structure formation is facilitated via a branching pi-stacks and/or hydrogen bonding. Interestingly, the glycosidic sugar groups can significantly influence the orientation of the secondary structure due to glycosidic backbone formation.

In our study of flavonoid behavior, the presence of additional hydroxyl groups within the glycosides and the flexibility associated with these sugars compared to the polyphenol core seem to decrease the probability of forming the most compact structures with the smallest RGYR. This reduced likelihood to form compact structures may indicate an ability to develop more diverse higher-dimensional structures that are larger and less compact. This distinction is notable when more complex flavonoids are compared to quercetin, which only has hydroxyl groups directly attached to the polyphenol core. The ability to form more complex structures would suggest a potential for building elaborate hierarchical arrangements with branching patterns, resulting in improved stabilization effects on both single and multiple macromolecules. Moreover, the flexible capacity of these flavonoids to form three-dimensional structures can likely facilitate adaptive interactions with other biomolecules, such as proteins, as we observed (**Fig. 2**). Our results are consistent with the possibility that the increased probability of developing flavonoid-mediated structures with higher dimensionality may contribute to enhanced stabilizing effects. Our studies suggest that flavonoid glycosidic moieties influence the assembly process, the diversity of fully assembled molecular states, and the formation of secondary structures.

Flavonoids, naturally upregulated in stressed plants, can protect against ultraviolet UV radiation^4,22^ and IDPs have been recently shown to protect organisms against environmental stressors like desiccation by forming fibers^14,23^. Thus, we investigated whether the ability of flavonoids to form supramolecular structures contributes to this protection by assessing the ability of quercetin, quercitrin, and isoquercitrin to protect cultured human fibroblasts from exposure to UV radiation. Treatment with these flavonoids resulted in concentration-dependent protection against the loss of cell viability caused by UV radiation, aligning with our previous studies (**Fig. 4a**). Notably, flavonoids with glycosidic sugar groups demonstrated the most robust protective effects, supporting our MDS finding that showed enhanced assembly of multidimensional structures by flavonoids containing sugar moieties.

**Fig. 4:**
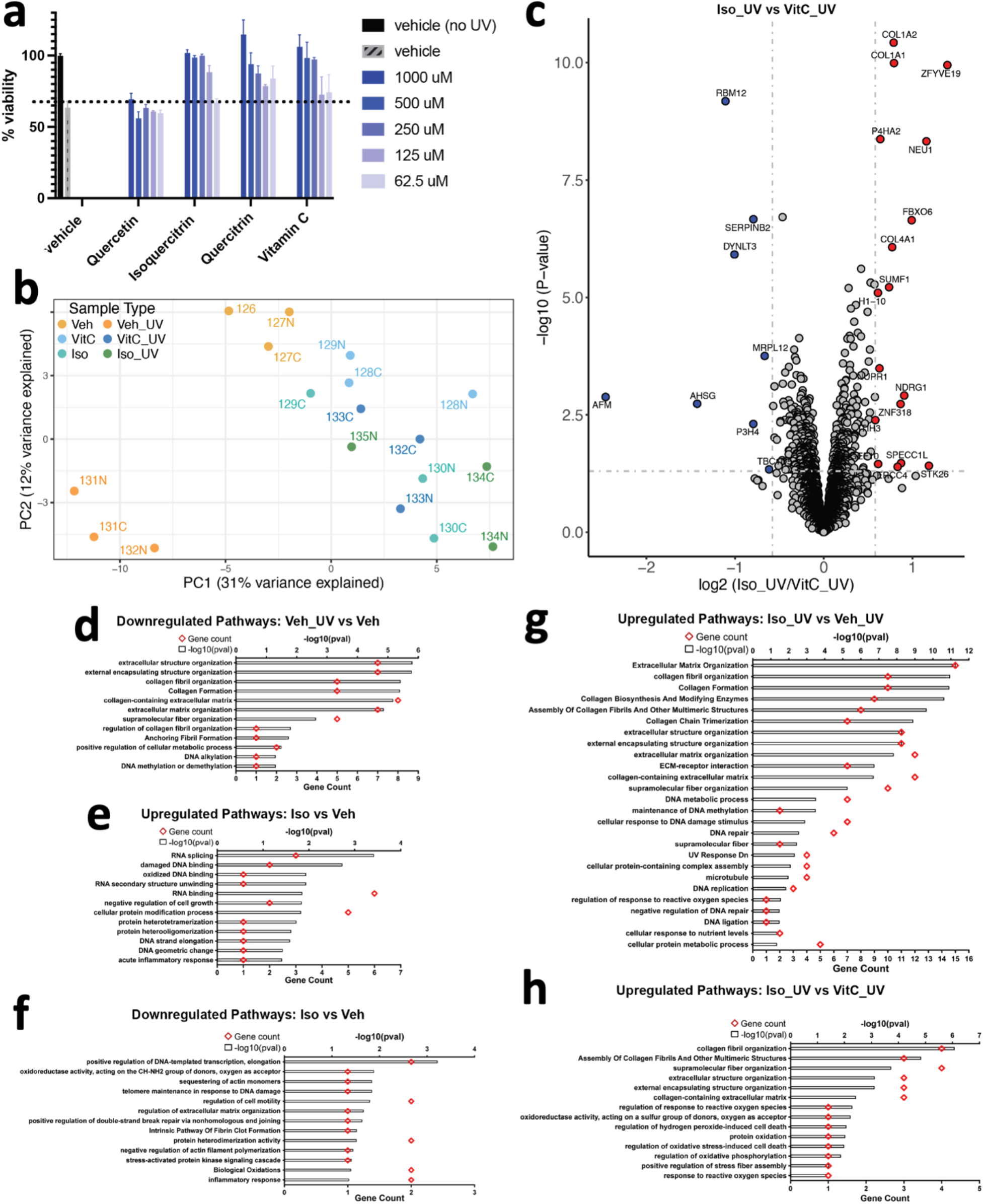
Flavonoids protect human dermal fibroblasts against ultraviolet radiation-induced cell death in a dose-dependent manner. **a**, Human fibroblasts were pretreated with the indicated concentration of compounds for 1h prior to ultraviolet radiation exposure. Cells were cultured in complete medium for an additional 24h, and cell viability was tested by CellTiter-Glo assay. Results are expressed as means ± SEM of three independent experiments. **b,** Principal Component Analysis (PCA) plot of proteomics data. Each data point represents a sample labeled with TMT (Tandem Mass Tag) and is colored according to its sample type. **c,** Volcano plot of proteins differentially expressed between fibroblasts exposed to UV light and treated with isoquercitrin or Vitamin C. Grey dotted horizontal and vertical lines denote a p-value cutoff of 0.05 and log2fc cutoff of 0.58 (1.5-fold change), respectively. Proteins denoted by red and blue dots are significantly up-and downregulated, respectively, in Isoquercitrin_UV samples relative to VitaminC_UV samples. **d,** Pathways downregulated in human fibroblasts upon UV exposure, in the absence of any treatment. **e-f,** Pathways up-and downregulated in isoquercitriin-treated fibroblasts relative to vitamin-C-treated fibroblasts, respectively, in the absence of UV exposure. **g-h,** Pathways upregulated in Isoquercitrtn_UV samples relative to Vehicle_UV (no treatment) and VitaminC_UV samples.

Many of the protective effects of flavonoids have been previously attributed to antioxidant effects^2^. Antioxidant effects can result from multiple mechanisms^24^ and so we conducted a proteomics study comparing the effects of flavonoids and the well known antioxidant Vitamin C, which is a free radical scavenger,^25^ to understand the underlying protective pathways. Our PCA plot (**Fig. 4b**) shows that cells treated with either Vitamin C or the flavonoid isoquercitrin, align with the untreated control on the PC1 axis, distinct from UV-treated cells, which implies a protective effect. Additionally, a more focused PCA plot comparing only cells treated with Vitamin C or flavonoid shows clear delineation based on treatment type (**Extended Data** Fig 3a), indicating that the cell protective mechanisms activated by these two treatments are distinct. Indeed, differential expression analysis showed distinct protein expression patterns between cells pretreated with flavonoids or Vitamin C (**Fig. 4c**) before being exposed to UV light as well as between untreated cells with and without UV exposure (**Extended Data Fig. 3b**). UV radiation upregulated cell death/senescence/necroptosis pathways (**Extended Data Fig. 3c)**, and downregulated collagens and proteins associated with cell cycle/DNA synthesis/repair (**Fig. 4d** and **Extended Data Fig. 3b**) in the absence of flavonoid. Under control conditions without UV treatment, the flavonoid isoquercitrin upregulated proteins associated with DNA repair pathways and downregulated inflammatory pathways (**Fig. 4e,f** and **Extended Data Fig. 3d**), while Vitamin C showed opposite effects (**Extended Data Fig. 3e,f,g**). When exposed to UV radiation, the presence of isoquercitrin induced upregulation of collagen and cell cycle/DNA repair proteins while downregulating proteins associated with autophagy, necroptosis, ferroptosis, lysosomal degradation, and other pathways (**Fig. 3g,h** and **Extended Dat**a Fig. 4h,i), whereas UV irradiated cells treated with Vitamin C primarily upregulated proteins associated with cell cycle/DNA repair-related pathways (**Extended Data Fig. 3k**). Moreover, Vitamin C treatment resulted in lower expression of proteins involved in pathways associated with cellular response to reactive oxygen species, which can be generated by exposure to UV radiation. These included STK26, which is involved in the regulation of response to reactive oxygen species (GO:1901031, GO:0000302), response to hydrogen peroxide (GO:0042542), regulation of oxidative stress-induced cell death (GO:1903201), and SUMF1, which is involved in protein oxidation (GO:0018158) and oxidoreductase activity (GO:0016670)^26–29^.

Notably, a comparison between isoquercitrin and Vitamin C in UV-irradiated cells revealed differential expression of various types of structural proteins (e.g., extracellular matrix, collagen, anchoring fibrils, etc.) and specifically, proteins involved in the supramolecular fiber organization pathway (GO:0097435) ^26,27^ (**Fig. 4h**). These results suggest that isoquercitrin may prevent the degradation of proteins found within regulatory pathways associated with higher-order structures, which is consistent with our models that predict isoquercitrin itself physically forms supramolecular structures and can interact physically with multiple proteins. These differences in protein expression patterns displayed by UV-irradiated cells treated with isoquercitrin versus Vitamin C raises the possibility that isoquercitrin enables cellular resilience to UV radiation by preventing the degradation of these multimolecular structural assemblies. Most importantly, our findings demonstrate that flavonoids and Vitamin C protect against UV radiation through different mechanisms and emphasize the role of flavonoids in supporting higher-order structural assemblies in a cellular context.

## Discussion

Flavonoids, commonly found in plant-based foods, exhibit remarkable abilities to regulate diverse biochemical activities and protect against cellular damage; however, the underlying mechanism is unknown. In this study, we discovered that flavonoids self-assemble into highly ordered supramolecular structures that influence enzymatic protein conformation, motion, and biochemical actions. Moreover, flavonoids with glycosidic groups, such as quercitrin and isoquercitrin, further influence this assembly process. Our findings also demonstrate that flavonoid assemblies play a role in the expression of enzyme structural protein assemblies under environmental stresses that provide increased cell resilience, preventing damage in response to UV radiation exposure. By uncovering this unique mechanism of flavonoid supramolecular assembly, this study opens a new path for understanding molecular and cellular regulation and provides new insight into the potential use of flavonoids as resilience therapeutics.

## Materials and methods

### Enzyme Assays

Enzyme screens were performed by SAMDI Tech (contract research organization); enzyme assays were performed in 20 µL volume in 384-well low-volume polypropylene microtiter plates (Greiner Bio-One) at room temperature. The enzymes (Arginase 1 (ARG1), Lysine demethylase 4C (KDM4C), MAP/microtubule affinity-regulating kinase 4 (MARK4), Histone-Lysine N-methyltransferase (NSD2), Tyrosine-protein phosphatase non-receptor type 1 (PTP1B), and Sirtuin-3 (SIRT3)) were incubated with compounds for 30 min prior to initiation of the reaction by the addition of substrate and cofactors (if required). Reactions were quenched by the addition of 0.5% formic acid (final) with subsequent neutralization using 1% sodium bicarbonate (final). For SAMDI MS analysis, 2 µL of each reaction was transferred using a 384-channel automated liquid handler to SAMDI biochip arrays functionalized with a neutravidin-presenting self-assembled monolayer. The preparation of SAMDI biochip arrays has been previously described ^30–32^. The SAMDI arrays were incubated for 1 h in a humified chamber to allow specific immobilization of the biotinylated substrates and products; they were then purified by washing with ultrafiltered water (50 µL/spot) and dried with compressed air. A matrix comprising α-Cyano-4-hydroxycinnamic acid in 80% acetonitrile: 20% aqueous ammonium citrate (10 mg/mL final) was applied in an automated format by dispensing 50 nL to each spot in the array. SAMDI MS was performed using reflector positive mode on an AB Sciex TOF-TOF 5800 System (AB Sciex, Framingham, MA) with 400 shots/spot analyzed in a random raster sampling. For data analysis, area under the curves (AUCs) for the product peak and substrate peak were calculated using the TOF/TOF Series Explorer (AB Sciex), and the amount of product formed was calculated using the equation AUC product / (AUC product + AUC Substrate). Negative controls were pre-quenched with 0.5% formic acid (final). Lysozyme assay was performed using the manufacturer’s instructions (Biovision Lysozyme inhibitor screening kit K237). Flavonoids were acquired from Sigma-Aldrich and Seleckchem.

### Enzyme assay analysis

*IC50 values of enzyme inhibition.* Inhibition of the enzymes was measured at 10 concentration values in the range of 5 nM to 100 µM for each compound and enzyme with two technical replicates, the percent inhibition values were determined by SAMDI software. Sigmoidal curves were fitted to the percent inhibition *vs* log_10_(*concentration*) series, and IC50 values were determined at the crossing point of 50 % inhibition. Manual verification of the fits and the IC50 values was performed. The resulting IC50 values fell into the following categories: a) within the assayed range of 5nM to 100µM (26 % of fits), b) IC50 could be extrapolated, IC50 > 100µM (12 % of fits), or c) fitting was not possible because all values were at baseline (62 % of fits). *Compound efficacy ranking*. First, for each IC50 value per compound and enzyme, a Z-score was calculated based on the log_10_(*IC50*) value and that of the compounds that produced a successful readout for enzyme *i*, according to eq 1:

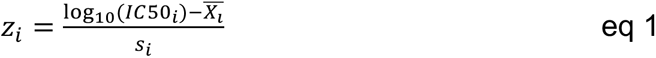

Where *X̅*_*i*_ and *s*_*i*_ are the average of all successfully determined log_10_(*IC50*) values for enzyme *i*. Any compound that did not produce a readout was assigned *z* = 2, this imputation allows ranking of compounds with very high IC50 values. Then, for each compound, a weighted mean of the z-scores (*z̅*) was calculated according to eq 2:

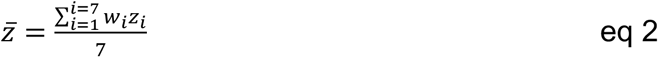

where the weights were calculated according to eq 3:

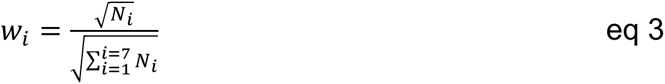

Where *N*_*i*_ is the number of successful readouts for enzyme *i*. Finally, for those compounds that had multiple repeated readouts, the average of the *z̅* values was used. Compounds were ranked based on the *z̅* values: low *z̅* values mean high efficacy of inhibition, and high values mean low efficacy. Fitting was performed using R (v3.6.3) and the gslnls package (v1.1.1), and figures were rendered using the ggplot2 package (v3.4.0).

### Molecular Dynamics Simulation

Initial protein structures were obtained from Uniprot AlphaFoldDB for MARK4 (AF-Q96L34-F1) and PTP1B (AF-P18031-F1), and the Lysozyme structure was acquired from the PDB (accession: 193L). Flavonoid structures were acquired from pubchem (quercetin CID-5280343, isoquercitrin CID-5280804, and quercitrin CID-5280459). Initial state input files for simulation were generated using AmberTools^33^. Amber forcefields were used and simulations were carried out with OpenMM^34^. RMSD and RGYR analysis were performed using the Python package mdtraj^35^. Visualizations of protein structures were produced using Houdini and the Python packages prody^36^, mdtraj, and biopython^37^. Weighted ensemble simulations were carried out using WEPY with 50 walkers, 1000 cycles, 10 ps steps, the REVOResampler, and an RGYR distance function. Final walker lineages and probability distribution were calculated based on the Lotz *et al.* pipeline and an RGYR distance function^20^.

### Cell culture and ultraviolet radiation

Human fibroblasts were obtained from Cell Applications Inc (Cat. # 106-05a) and were maintained in Dulbecco’s Modified Eagle medium (DMEM, Corning, Cat. #10-013-CV) with 10% fetal bovine serum (Gibco, Cat. #10082147) and 1% penicillin/streptomycin (Gibco, Cat. # 15070-063) in a humidified incubator with 5% CO_2_ at 37 °C. For ultraviolet radiation, fibroblasts were seeded in a 96-well plate at a density of 25 000 cells/well and maintained at 37 °C for 24 h. Cells were washed once with Hanks’ balanced salt solution (HBSS, Gibco, Cat. #14025-092), then treated for 1 h with different concentrations of compounds prepared in HBSS. Vitamin C (Sigma, Cat. # A5960) served as a positive control. Treated cells were exposed to ultraviolet light (UVP cross-linker, CL1000, 254 nm UV, 8 watts) at a total dose of 20 mJ/cm^2^. Following irradiation, the HBSS was replaced with culture medium containing the same concentration of compounds as before irradiation, and cells were again maintained at 37 °C. Cell viability was measured at 24 h post-irradiation using CellTiter-Glo according to the manufacturer’s instructions (Promega, Cat.# G7571). Results were expressed as relative cell viability (%) with respect to control cells (cells without ultraviolet irradiation or compound treatment).

### Proteomics sample preparation

Frozen cell pellets were dissolved for 15 min in 6M Guanidinium Chloride at room temperature to lyse cells. Then each solution was run on a 10 kDa filter ( Pall, TX) and washed with 50 µL of TEAB buffer at 3000 rpm. **FASP Digestion** ^39^: A stock solution of Trypsin Platinum, Mass Spec Grade (Promega) at 100 µg/mL in 50 mM TEAB, and a specific volume of stock trypsin was added to each sample vial for each set of brain cells using a 1:50 ratio (Trypsin:Protein). Samples were incubated for 2 h at 50 °C and shaken/mixed at 350 rpm on an Eppendorf ThermoMixer C. **TMT Labeling:** Samples, now peptides, were labeled using 4 µL of TMTpro Mass Tag Labels (ThermoScientific). After labeling, samples were then shaken/mixed for 45 min to ensure labels were covalently bonded to peptides. The labeling reaction was quenched for 10 min using 1 µL of 5% hydroxyalamine. After quenching, samples were pooled into one 2 mL Eppendorf tube and then dried down using an Eppendorf Vacufuge Plus. **Desalting Samples:** Dried samples were resuspended in 300 µL of 0.1% TFA in ultrapure HPLC grade water and vortexed to ensure full solubility. Samples were then desalted using Pierce^TM^ Peptide Desalting Spin Columns (ThermoScientific). Final eluates (desalted samples) contained 600 µL of 50% HPLC grade Acetonitrile and 50% ultrapure HPLC grade water. Desalted samples were then transferred to HPLC vials and were again dried down. Dried and desalted samples were resuspended in 6 µL of 0.1% Formic Acid in ultrapure HPLC grade water and were then injected for LC-MS/MS analysis.

### Mass spectrometry analysis

After separation, each fraction was submitted for a single LC-MS/MS experiment performed on an HFX Orbitrap (Thermo Scientific) equipped with an Exigent (SCIEX, MA) nanoHPLC pump. Peptides were separated onto a 150 µm x 8 cm PepSep C18 analytical column (Bruker, MA) by applying a gradient from 5–25% ACN in 0.1% formic acid over 90 min at 300 nL min−1. Electrospray ionization was enabled by applying a voltage of 2 kV using a PepSep electrode junction at the end of the analytical column and sprayed from stainless steel PepSep emitter SS 30 µm LJ (Odense, Denmark). The HF Orbitrap was operated in data-dependent mode for the mass spectrometry methods. The mass spectrometry survey scan was performed in the Orbitrap in the range of 450 –900 m/z at a resolution of 1.2 × 10^5^, followed by selection of the ten most intense ions. The (TOP10) ions were then subjected to an HCD MS2 event in the Orbitrap part of the instrument. The fragment ion isolation width was set to 0.8 m/z, AGC was set to 50,000, the maximum ion time was 150 ms, normalized collision energy was set to 34V, and an activation time of 1 ms was set for each HCD MS2 scan.

### Mass spectrometry data analysis

Raw data were submitted for analysis in Proteome Discoverer 3.0.1.23 (Thermo Scientific) software with Chimerys. Assignment of MS/MS spectra was performed using the Sequest HT algorithm and Chimerys (MSAID, Germany) by searching the data against a protein sequence database including all entries from the Mouse Uniprot database (SwissProt 19,768 2019) and other known contaminants such as human keratins and common lab contaminants. Sequest HT searches were performed using a 20 ppm precursor ion tolerance and requirement of each peptide N-/C termini to adhere with Trypsin protease specificity while allowing up to two missed cleavages. 18-plex TMT tags on peptide N termini and lysine residues (+304.207146 Da) were set as static modifications, and Carbamidomethyl on cysteine amino acids (+57.021464 Da), while methionine oxidation (+15.99492 Da) was set as a variable modification. An MS2 spectra assignment false discovery rate (FDR) of 1% on the protein level was achieved by applying the target-decoy database search. Filtering was performed using a Percolator (64-bit version, (*11*)). For quantification, a 0.02 m/z window centered on the theoretical m/z value of each of the six reporter ions, and the intensity of the signal closest to the theoretical m/z value was recorded. Reporter ion intensities were exported to the result file of the Proteome Discoverer 3.0 search engine as Excel tables. The total signal intensity across all peptides quantified was summed for each TMT channel, and all intensity values were adjusted to account for potentially uneven TMT labeling and/or sample handling variance for each labeled channel.

### Proteomics analysis

Protein level data from each TMT channel was analyzed in R. The dataset was first preprocessed by removing contaminant proteins and those with unavailable abundance data. Next, the dataset was normalized using the variance stabilizing normalization method (vsn) with the R MSnbase package. Differential expression (DE) analysis utilized the R limma package with volcano plots for visualization (pval ≤ 0.05, log2fc ≥ |0.58| for significance). Only the proteins that were identified as significantly differentially expressed were selected for further pathway analysis, which was performed with the R Enrichr package, identifying significantly altered pathways (pval ≤ 0.05) for up-and downregulated proteins in each DE comparison.

## Notes

### Competing Interest Statement

The authors have declared no competing interest.

