## Supplementary material for "Control of Molecular Biochemistry and Cell Injury Responses through Highly Ordered Supramolecular Assembly of Flavonoids": SI Appendix

| name | Uniprot ID | Enzyme | structure | description |
|---|---|---|---|---|
| **MARK4** | Q96L34 | Microtubule affinity regulating kinase 4 | monomeric | serine/threonine phosphorylation |
| **KDM4C** | Q9H3R0 | Lysine demethylase 4C | monomeric | demethylation of histones |
| **Lysozyme** | P00698 | Lysozyme | monomeric | Breaks glycosidic bonds |
| **NSD2** | O96028 | Histone-Lysine N-methyltransferase | monomeric | Epigenetic control of gene expression |
| **PTP1B** | P18031 | Protein tyrosine phosphatase | monomeric | Negative regulator of insulin signaling pathway. |
| **ARG1** | P05089 | Arginase I | trimeric | Converts arginine to ornithine and urea |
| **SIRT3** | Q9NTG7 | Siturin-3 | dimeric | Mitochondrial NAD-dependent deacetylase |

**Extended Data Table. 1: Enzymes used for biochemical screens.**

Summary table of compound ranking and enzymatic activity assays

| Name | Structure | Rank | ARG1 | KDM4C | Lysozyme | MARK4 | NSD2 | PTP1B | SIRT3 |
|---|---|---|---|---|---|---|---|---|---|
| *quercetin* | | 12.14 | | | | | | | |
| *isoquercitrin* | | 49.86 | | | | | | | |
| *orientin* | | 37.43 | | | | | | | |
| *mangiferin* | | 43.71 | | | | | | | |
| *gossypin* | | 46.57 | | | | | | | |
| *tiliroside* | | 53.14 | | | | | | | |

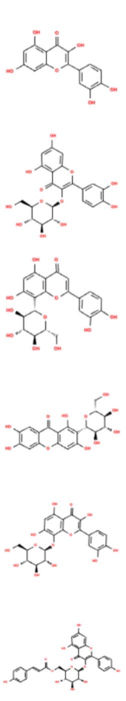

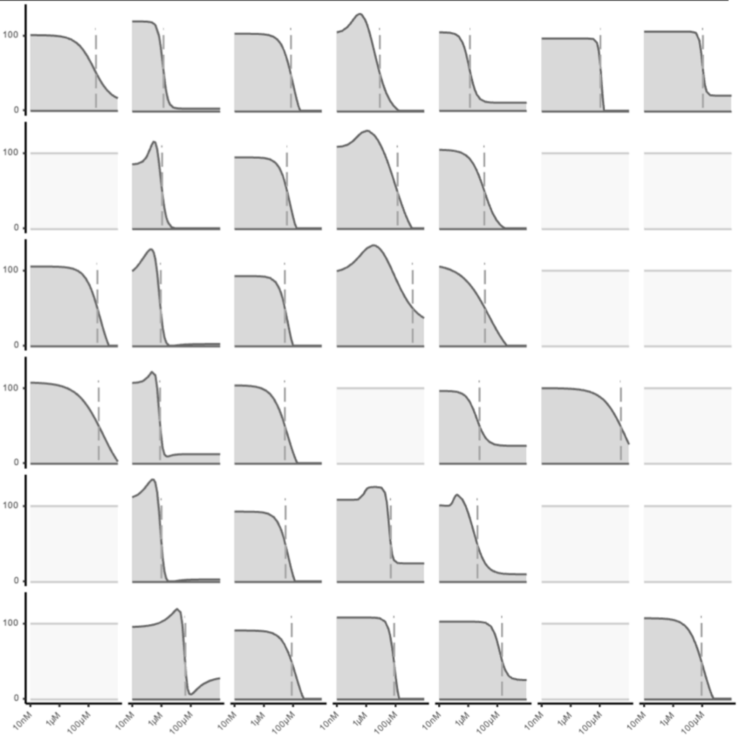


**Extended Data Table. 2a.**

Summary table of compound ranking and enzymatic activity assays

| Name | Structure | Rank |
| --- | --- | --- |
| *leucoside* | 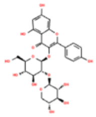 | 54.86 |
| *myricitrin* | 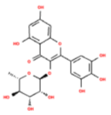 | 58.86 |
| *rutin* | 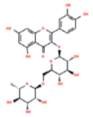 | 59.14 |
| *guaijaverin* | 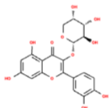 | 59.86 |
| *rhamnetin 3-sophoroside* | 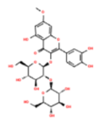 | 61.43 |
| *avicularin* | 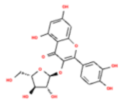 | 63.43 |

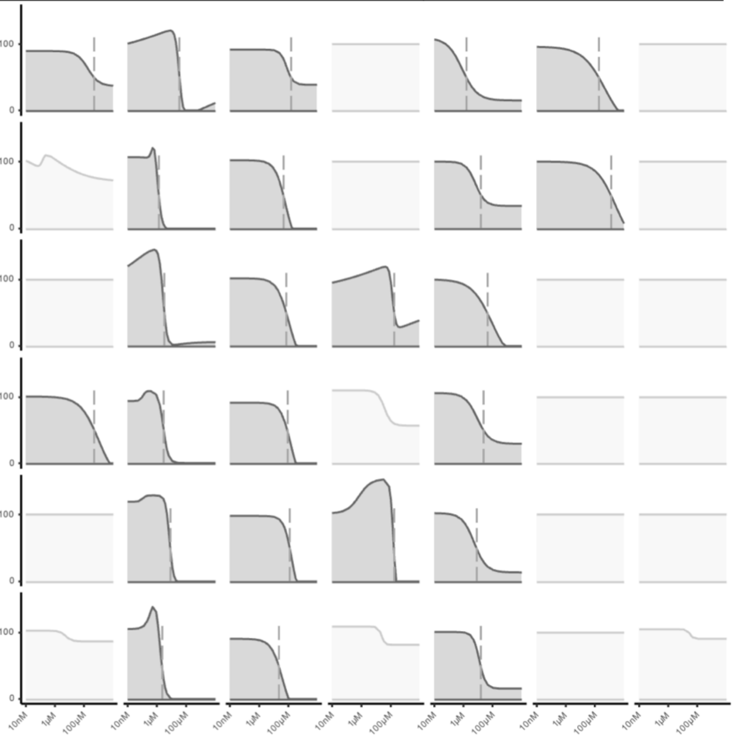


**Extended Data Table. 2b.**

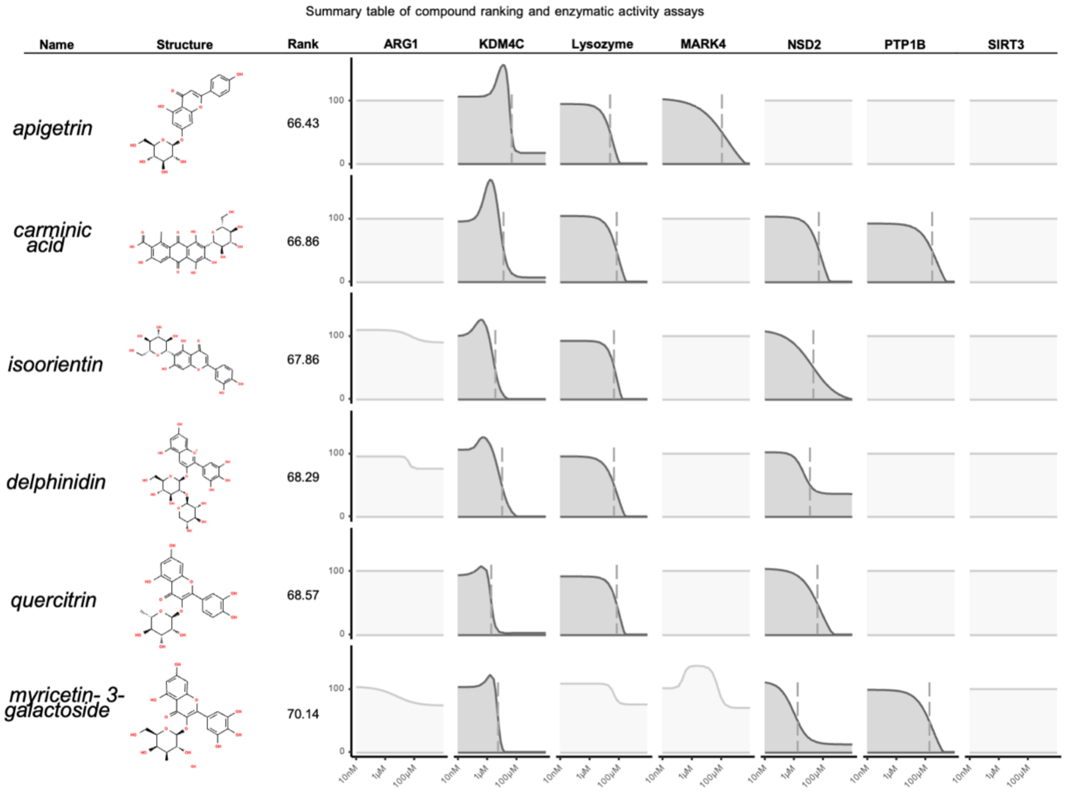


**Extended Data Table. 2c.**

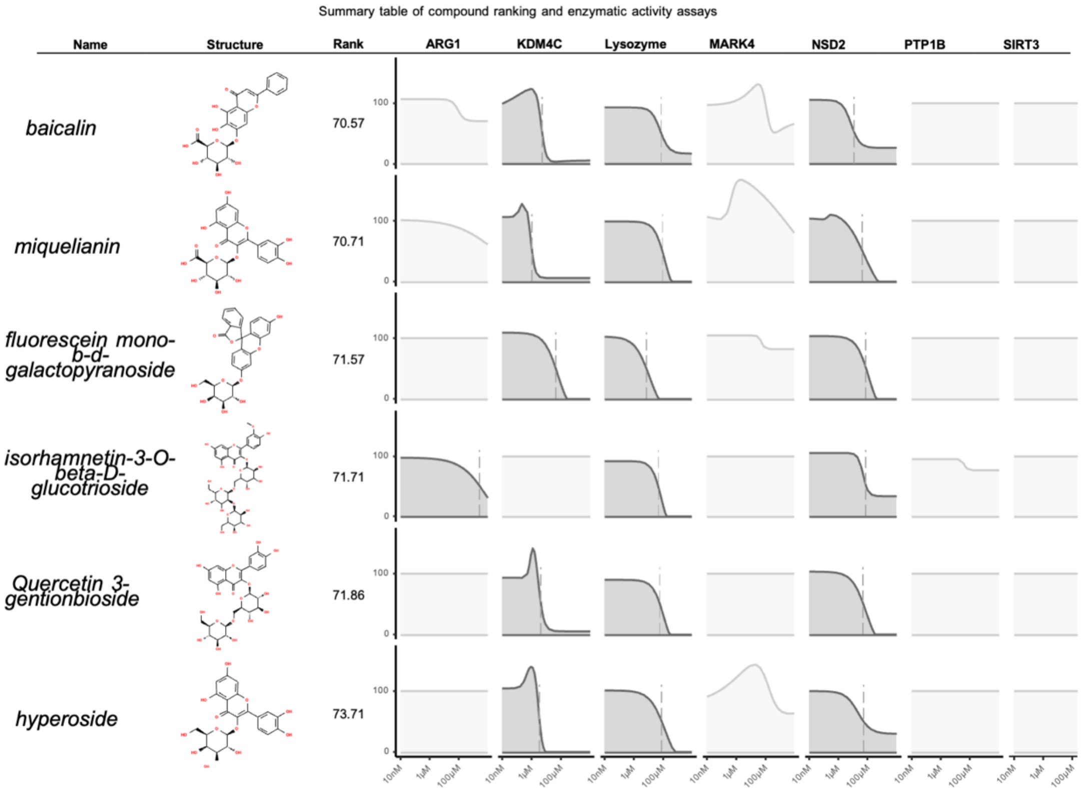


**Extended Data Table.** 2d.

Summary table of compound ranking and enzymatic activity assays

| Name | Structure | Rank |
|---|---|---|
| *Quercetin 3,4'-diglucoside* | 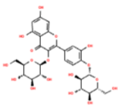 | 73.71 |
| *spiraeoside* | 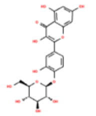 | 76.43 |
| *quercetin 3-o-malonylglucoside* | 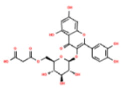 | 78.29 |
| *haploside C* | 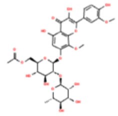 | 79 |
| *mearnsitrin* | 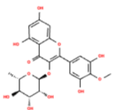 | 79 |
| *Peonidin-3-O-rutinoside chloride* | 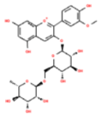 | 79.57 |

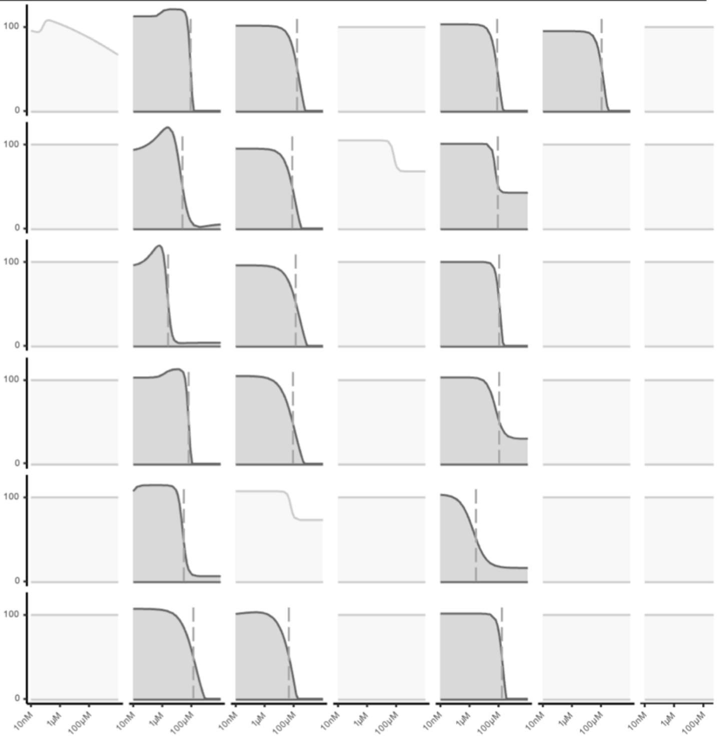


**Extended Data Table.** 2e.

Summary table of compound ranking and enzymatic activity assays

| Name | Structure | Rank |
|---|---|---|
| *astragalin* | 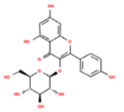 | 79.86 |
| *isoliquiritin apioside* | 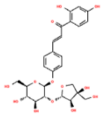 | 80 |
| *pectolinarin* | 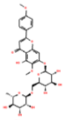 | 80.14 |
| *epimedin B* | 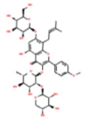 | 81.14 |
| *epimedin A* | 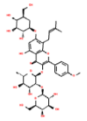 | 81.29 |
| *kaempferol 7-O-neohesperidoside* | 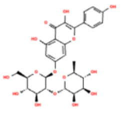 | 82.71 |

ARG1 | KDM4C | Lysozyme | MARK4 | NSD2 | PTP1B | SIRT3

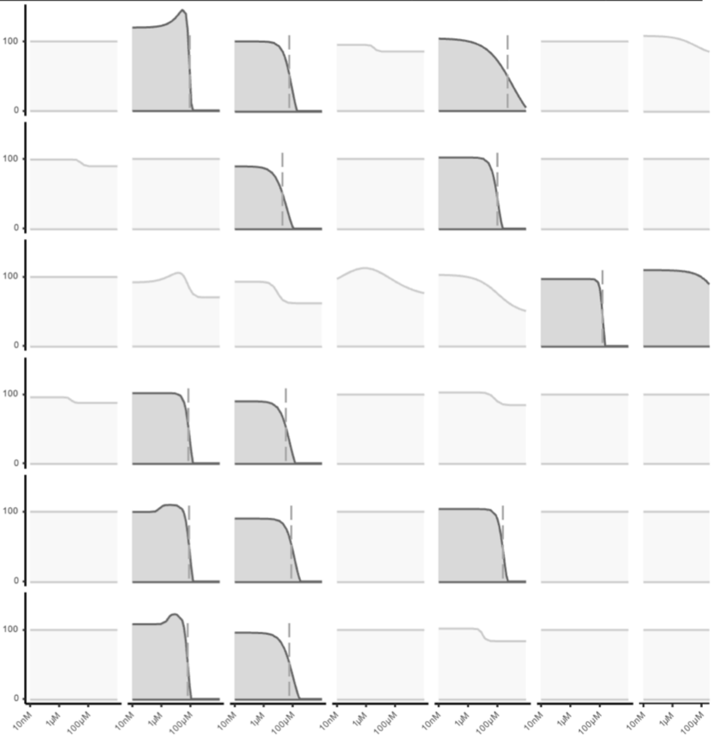


**Extended Data Table. 2f.**

Summary table of compound ranking and enzymatic activity assays

| Name | Structure | Rank |
|---|---|---|
| *oroxindin* | 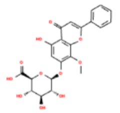 | 84 |
| *Apigenin 7-O-(2G-rhamnosyl) gentiobioside* | 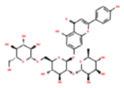 | 85 |
| *narcissin* | 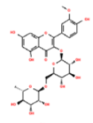 | 85.71 |
| *calendoflavoside* | 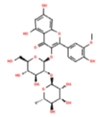 | 86.43 |
| *naringin* | 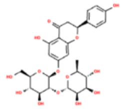 | 87.29 |
| *typhaneoside* | 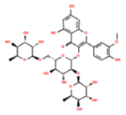 | 87.57 |

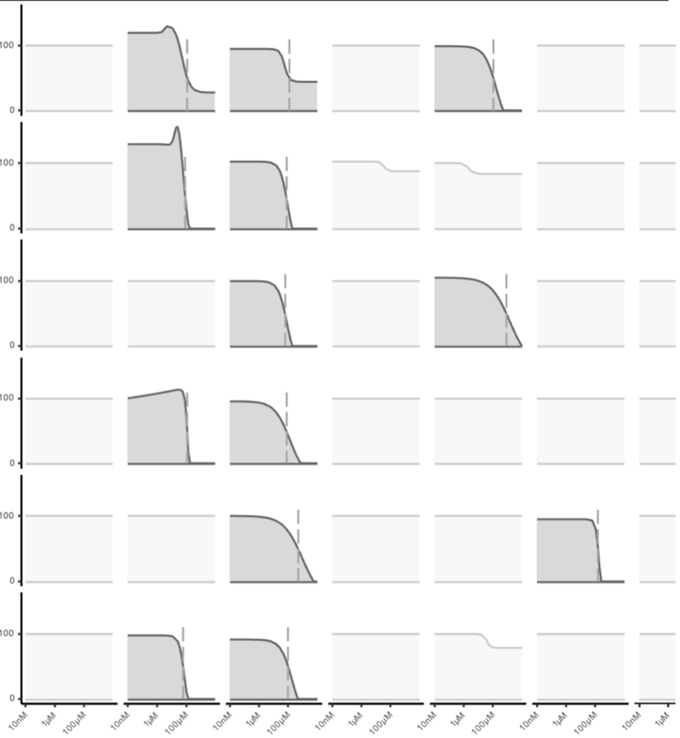


**Extended Data Table. 2g.**

Summary table of compound ranking and enzymatic activity assays

| Name | Structure | Rank |
|---|---|---|
| *robinin* | 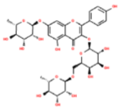 | 87.86 |
| *vitexin* | 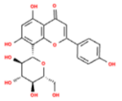 | 89.29 |
| *icariin* | 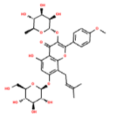 | 89.43 |
| *kaempferol 3-rutinoside 4'-glucoside* | 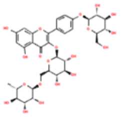 | 90.57 |
| *kaempferitrin* | 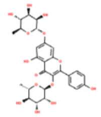 | 91 |
| *afzelin* | 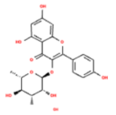 | 92.29 |

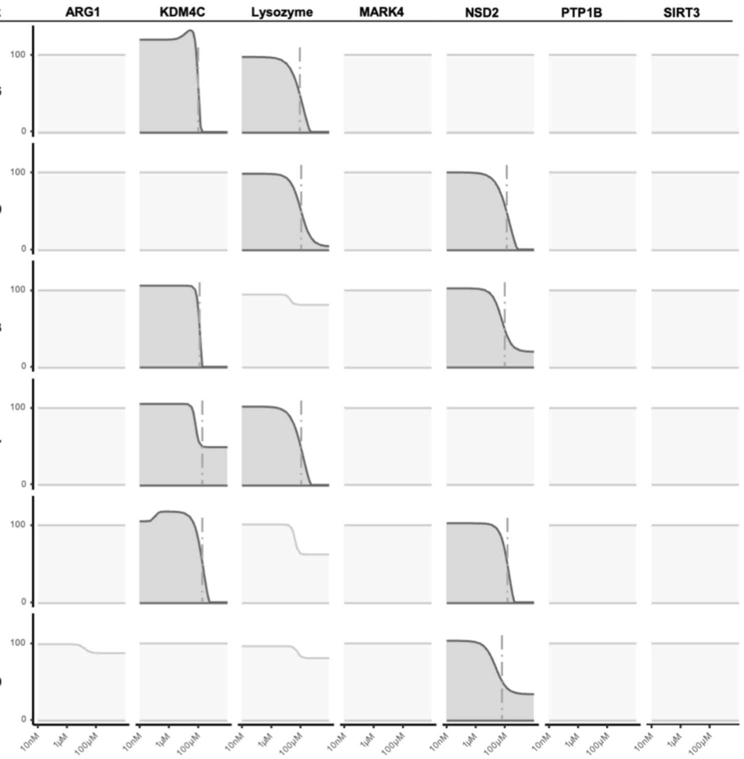


**Extended Data Table. 2h.**

Summary table of compound ranking and enzymatic activity assays

| Name | Structure | Rank | ARG1 | KDM4C | Lysozyme | MARK4 | NSD2 | PTP1B | SIRT3 |
|---|---|---|---|---|---|---|---|---|---|
| *Vitexin 2-O-rhamnoside* | 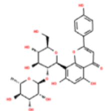 | 93 | | | | | | | |
| *epimedin C* | 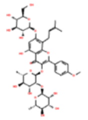 | 94 | | | | | | | |
| *populin* | 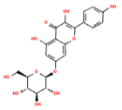 | 94 | | | | | | | |
| *Naringin dihydrochalcone* | 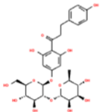 | 95.29 | | | | | | | |
| *nicotiflorin* | 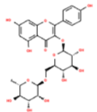 | 95.71 | | | | | | | |
| *kaempferol 3-gentiobioside* | 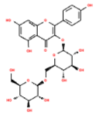 | 96.29 | | | | | | | |

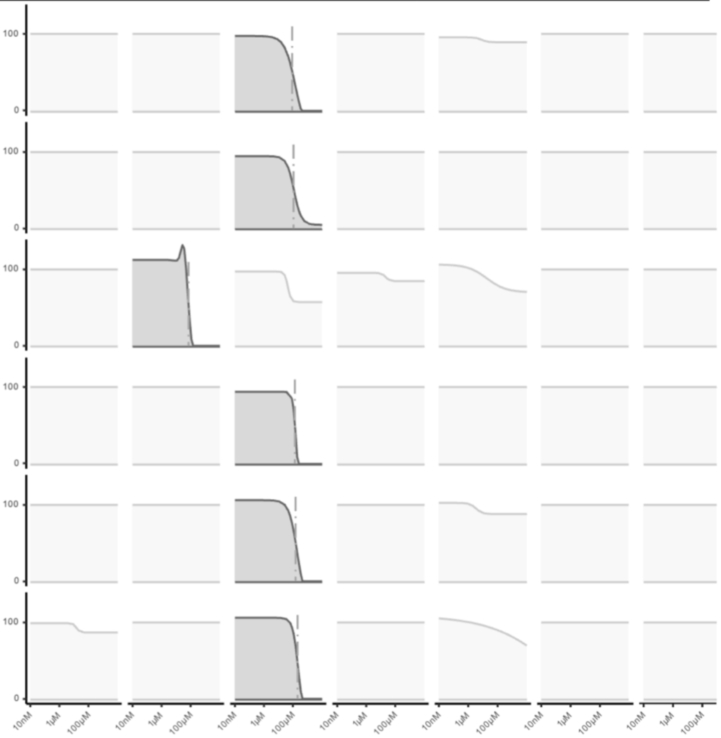


**Extended Data Table. 2i.**

Summary table of compound ranking and enzymatic activity assays

| Name | Structure | Rank |
|---|---|---|
| chromen-2-one xylofuranoside | 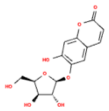 | 96.57 |
| *fluorescein-digalactoside* | 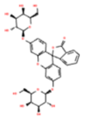 | 96.71 |
| *camelliaside* | 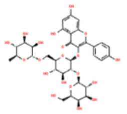 | 100 |
| *troxerutin* | 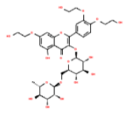 | 100 |

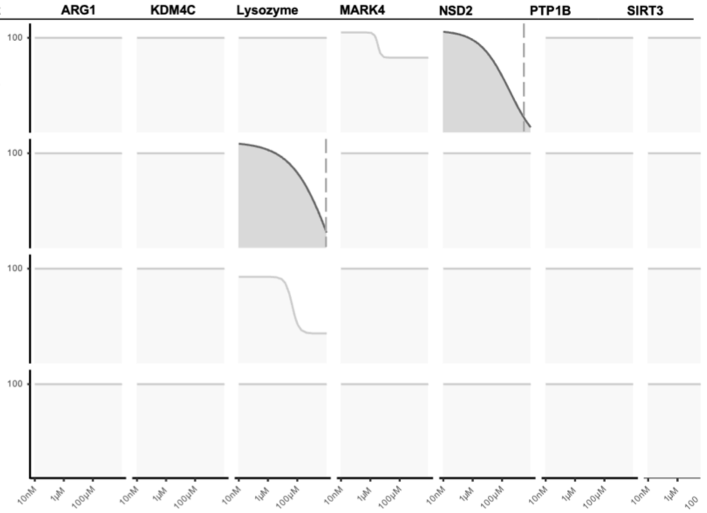


**Extended Data Table. 2j.**

**Extended Data Table. 2: Dose-dependent inhibition of enzyme activity by a library of flavonoids.**

**a-j**, Concentration-dependent inhibition observed for 58 flavonoids in enzyme activity assays for Arginase 1 (ARG1), Lysine demethylase 4C (KDM4C), Lysozyme, MAP/microtubule affinity-regulating kinase 4 (MARK4), Histone-Lysine N-methyltransferase (NSD2), Tyrosine-protein phosphatase non-receptor type 1 (PTP1B), and Siturin-3 (SIRT3). Compounds are sorted based on the decreased order of overall efficacy of enzyme inhibition; for the ranking methodology see the Materials and Methods section.

**Extended Data Table. 3: Results of differential expression and pathway analysis of proteomics data. [See .xlsx file]**

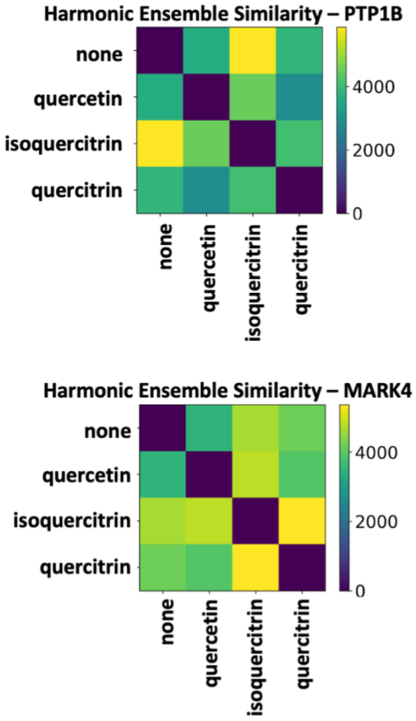


**Extended Data Fig.1: Analysis of MDS protein dynamics.**

Harmonic ensemble similarity analysis was performed on the last 10 ns of simulation in figure 2b, using PTP1B core enzyme backbone atoms for residues 3 to 277 (top) and MARK4 core enzyme backbone atoms for residues 59 to 310 (bottom) in the presence of flavonoids and alone.

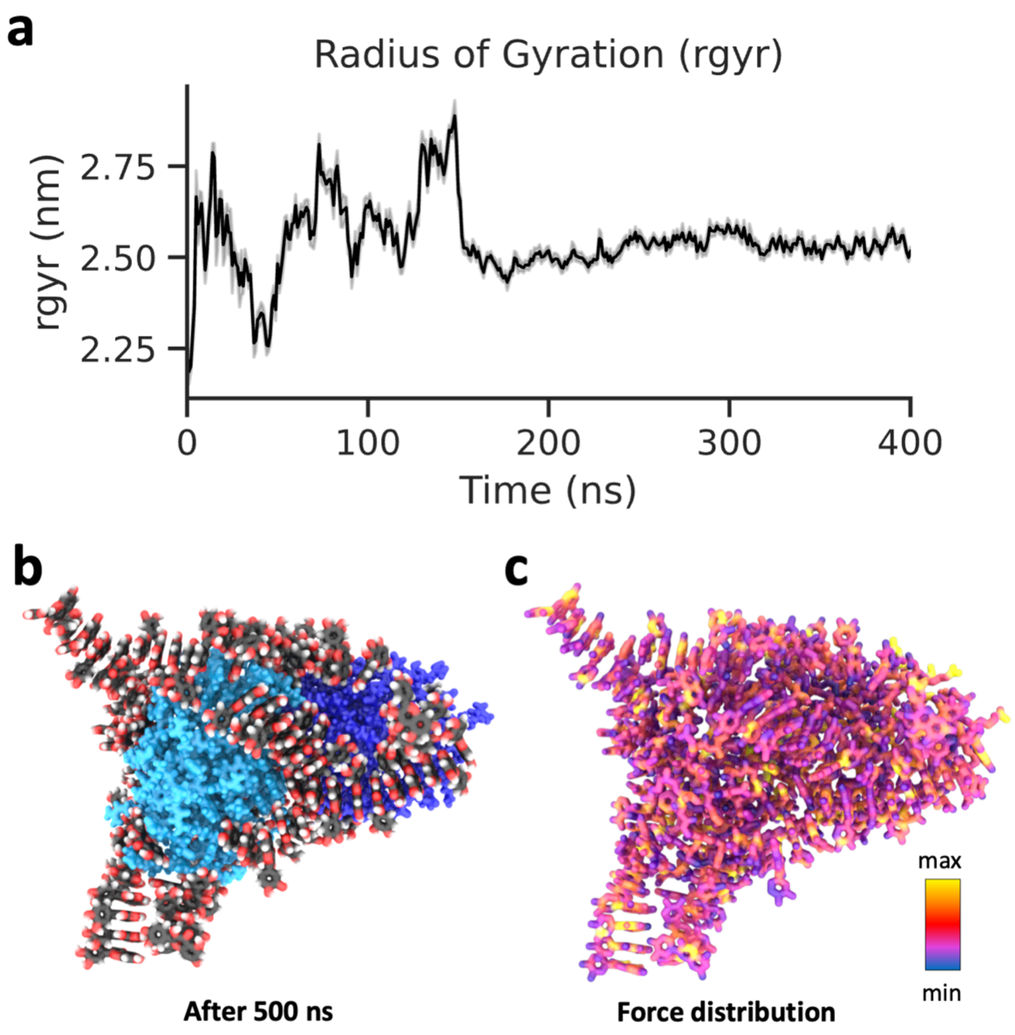


**Extended Data Fig. 2: MDS reveals flavonoid supramolecular structures influence intermolecular mobility and mechanical force distribution.**

**a,** Radius of gyration (RGYR) from MDS of two lysozyme molecules and 128 quercetin molecules. **b,** A single state from the end of the 400 ns MDS in **a**. **c**, The same state as **b** rendered to indicate the relative force experienced by each atom**.**

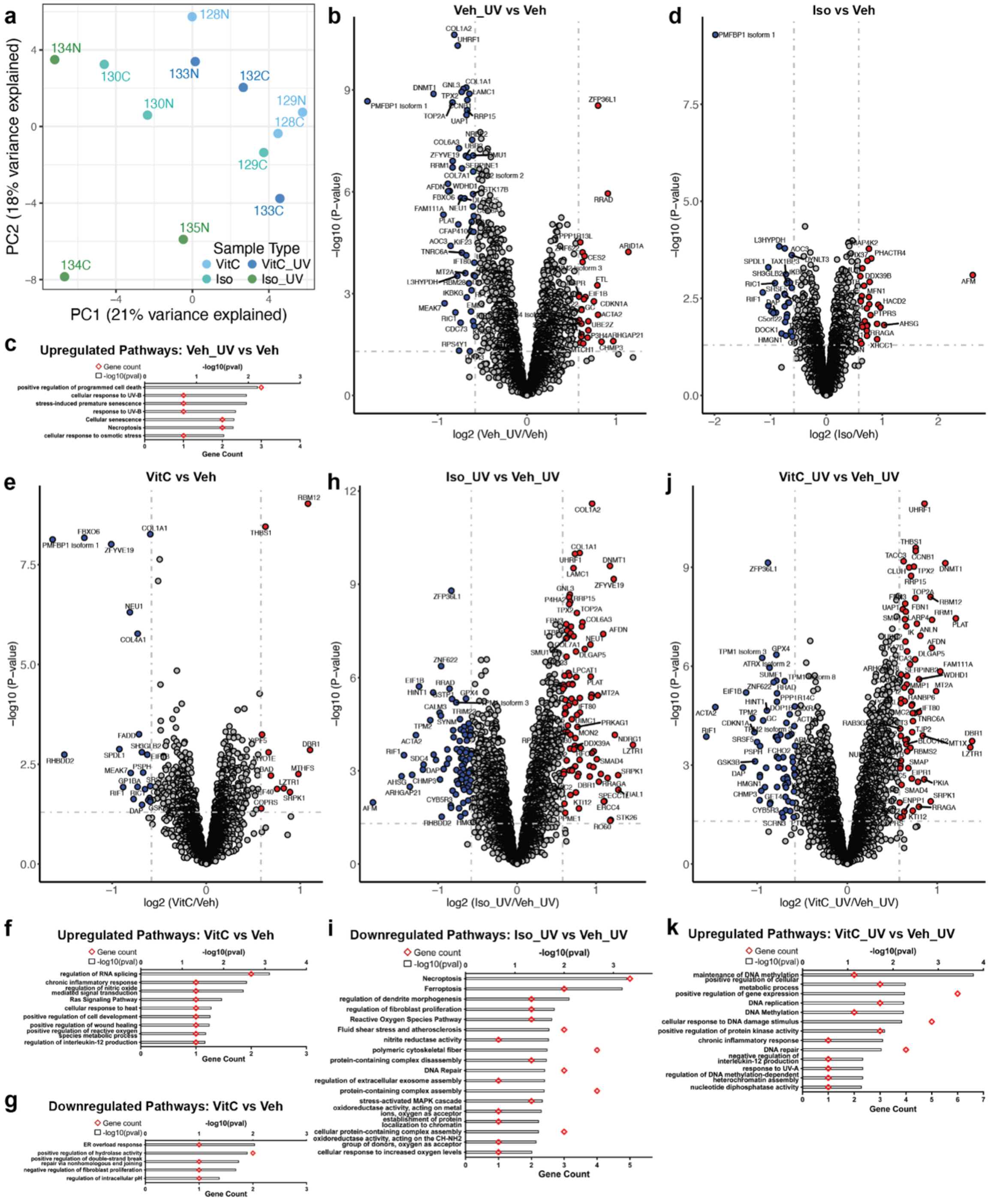


**Extended Data Fig. 3**

**Extended Data Fig. 3: Proteomics data analyses suggests treatment of UV-exposed fibroblasts with isoquercitrin and vitamin C induces protection through distinct mechanisms and pathways.**

**a,** Focused PCA plot showing clustering of only the proteomics samples treated with isoquercitrin or vitamin C, with and without UV-exposure. **b, d, e, h, j,** Volcano plots of proteins differentially expressed in Veh_UV vs Veh, Iso vs Veh, VitC vs Veh, Iso_UV vs Veh_UV and Vitc_UV vs Veh_UV proteomic profile comparisons, respectively. Grey dotted horizontal and vertical lines denote a p-value cutoff of 0.05 and log2fc cutoff of 0.58 (1.5-fold change), respectively. Proteins denoted by red and blue dots are significantly up- and downregulated, respectively. **c, f, k,** Pathways upregulated in Veh_UV vs Veh, VitC vs Veh, and VitC_UV vs Veh_UV proteomic profile comparisons, respectively. **g, i,** Pathways downregulated in VitC vs Veh and Iso_UV vs Veh_UV proteomic profile comparisons, respectively.
